# Nitric oxide suppresses cilia activity in ctenophores

**DOI:** 10.1101/2023.04.27.538508

**Authors:** Tigran P. Norekian, Leonid L. Moroz

**Author notes:** **Correspondence:** Leonid L. Moroz. Equal contribution.

## Abstract

Cilia are the major effectors in Ctenophores, but very little is known about their transmitter control and integration. Here, we present a simple protocol to monitor and quantify cilia activity in semi-intact preparations and provide evidence for polysynaptic control of cilia coordination in ctenophores. Next, we screen the effects of several classical bilaterian neurotransmitters (acetylcholine, dopamine, L-DOPA, serotonin, octopamine, histamine, gamma-aminobutyric acid (GABA), L-aspartate, L-glutamate, glycine), neuropeptides (FMRFamide), and nitric oxide (NO) on cilia beating in *Pleurobrachia bachei* and *Bolinopsis infundibulum*. Only NO inhibited cilia beating, whereas other tested transmitters were ineffective. These findings further suggest that ctenophore-specific neuropeptides could be major candidate signaling molecules controlling cilia activity in representatives of this early-branching metazoan lineage.

## INTRODUCTION

Ctenophores or comb jellies have the largest cilia in the animal kingdom, primarily used for complex locomotion in most species within this phylum. Moreover, cilia contribute to the majority of functions and behaviors of ctenophores (Tamm, 1982;Tamm, 2014). One primary example is ctene rows [or ctene plates], which consist of the large mechanically fused swim cilia used by ctenophores to move in the water column. The coordination of multiple behaviors in ctenophores is controlled by variations in the activity of swim cilia, and these mechanisms were under intensive investigation (Tamm and Tamm, 1981;Tamm, 1983;1984;Nakamura and Tamm, 1985;Moss and Tamm, 1986;1987;Tamm, 1988;Tamm and Tamm, 1988;Tamm and Terasaki, 1994).

Although cilia are the main effectors in ctenophores, with presumed neuronal control and different subtypes of synapses detected by electron microscopy (Hernandez-Nicaise, 1991;Burkhardt et al., 2023), little is known about synaptic regulation and neurotransmitters controlling cilia movement. Initial identification of glutamate as a small signal molecule and neurotransmitter candidate in ctenophores (Moroz et al., 2014;Moroz et al., 2020b;Moroz et al., 2021) targeted muscular systems. Still, in early experiments, glutamate did not change cilia beating (Moroz et al., 2014), and some ionotropic glutamate receptors were sensitive to glycine (Alberstein et al., 2015;Yu et al., 2016).

It was proposed that neural systems evolved independently in ctenophores by developing a unique molecular and structural organization (Moroz, 2014;Dabe et al., 2015;Kohn et al., 2015;Moroz, 2015;Moroz and Kohn, 2015;Whelan et al., 2015), including a subset of ctenophore-specific secretory peptides that could act as signal molecules (Moroz, 2014;Moroz et al., 2014;Moroz, 2021). Multiple candidates were identified in *Pleurobrachia* and *Mnemiopsis* (Moroz et al., 2014;Moroz and Kohn, 2016). The recent genome-wide and mass spectroscopy survey further expanded the list of secretory peptide candidates and identified some (neuro)peptides involved in the control of cilia beating in juvenile *Mnemiopsis* (Sachkova et al., 2021) and *Bolinopsis* (Hayakawa et al., 2022). However, cellular bases of ctenophore behavior are unknown.

Quantitative recording of cilia activity in ctenophores is equally essential for behavioral and functional analyses in both juvenile and adult animals. First, we describe a simple protocol successfully used to record and quantify cilia beating in ctenophores. This protocol can be practical for screening and investigating the physiological roles of different transmitters. Second, we provide initial evidence of (i) polysynaptic control of cilia coordination using chemical transmission, (ii) confirmed negative results of classical bilaterian neurotransmitter action on cilia, and (iii) showed a potential regulatory role of the gaseous molecule, nitric oxide (NO), in cilia beating.

## MATERIALS AND METHODS

Large, 1-to-2 cm, *Pleurobrachia bachei* and medium-size, 3-to-4 cm, *Bolinopsis infundibulum* were collected from the dock at Friday Harbor Laboratories, University of Washington, in the Pacific Northwest. Initially, freshly caught animals were incubated in high Mg^2+^ seawater (333 mM magnesium chloride added to filtered seawater at a 1:1 ratio) for about 15-60 minutes to prevent muscle contractions, especially for larger animals. The animals were then tightly pinned to a Sylgard-coated Petri dish (World Precision Instruments, Sylgard Silicone Elastomer, SYLG184) with small steel insect pins to prevent all body movements other than cilia beating. The excess of MgCl_2_ was washed out without any noticeable effect on cilia beating compared to control, and semi-intact preparations could maintain cilia activity within several hours. Small animals (<1 cm *Pleurobrachia*) were used whole without dissection; larger animals (2 cm and above) were dissected, and parts of a body wall with 2-3 cilia rows were pinned the same way to the Petri dish.

The Petri dish was placed in a standard electrophysiological rig on a recording platform and connected to the Ag/AgCl reference electrode. We used glass microelectrodes (borosilicate glass micropipettes for intracellular recording from World Precision Instruments - standard glass capillaries 2 mm diameter with a thin filament, 1B200F-4), filled with 3M potassium acetate to record cilia beating. The sharp microelectrodes were pulled using Microelectrode Puller (Sutter Instruments, Flaming/Brown Micropipette Puller P-97). The original resistance of sharp microelectrodes (made for intracellular recordings) was around 30 MΩ. A narrow strip of thin paper was used to carefully touch the tip of the electrode to break off the most fragile sharp end. The resulting electrode was more stable to further mechanical contact and had a resistance of 3-5 MΩ. Electrodes with very low resistances (below 1 MΩ) and wider tips were unsuitable. The electrodes were then connected to the micromanipulators (Warner Instruments, Standard Manual Control Micromanipulators, MM-33) and the intracellular amplifiers (Neuroprobe 1600, A-M Systems).

With the help of micromanipulators and under visual control via a dissecting microscope (Nikon stereoscopic microscope SMZ-10A), the tip of the electrode was carefully placed next to the cilia combs so that during cilia beating, cilia were touching the end of the electrode, providing electrical signals (**Figure 1A**). This physical contact created a brief electrical signal picked up by amplifiers and recorded on paper and in digital form using Gould Recorder (WindoGraf 980). Thus, each cilia beat was translated into a fast electrical spike. Combining electrophysiology with microscopy, we observed a one-to-one relationship between a cilia strike and a recorded electrical signal/spike, which allowed a digital recording of cilia beat frequency. It is important to note that this technique did not allow quantification of cilia beating amplitude and forces - only the frequency. It was crucial for stable recording to have the ctenophore body wall tightly pinned to the Sylgard-coated Petri dish, with no movements except cilia beating.

**Figure 1.**
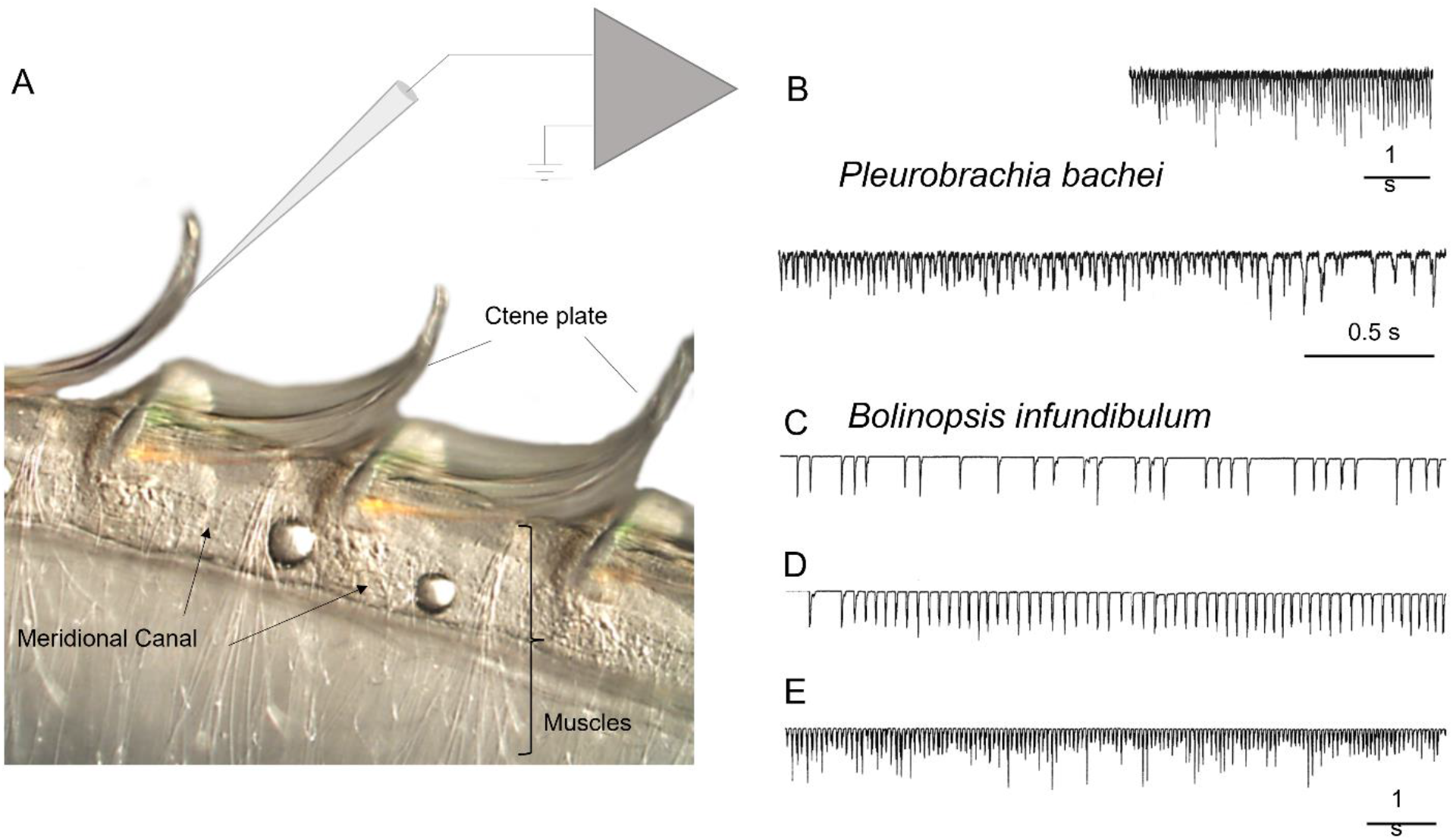
Recording cilia activity in *Bolinopsis* and *Pleurobrachia*. A. Schematic diagram showing the position of a recording microelectrode near comb plates in *Bolinopsis* (see text for details). Illustrative examples of cilia beating recording in *Pleurobrachia* (B) and *Bolinopsis* (C-E).

In both tested species, muscle contractions did not affect the recording of cilia in our semi-intact preparations. Thus, in most pharmacological tests, we did not apply elevated MgCl_2_. We used MgCl_2_ only to suppress chemical/synaptic transmission and excitability in particular experiments described in the results. In these experiments, applied 150-200 mM MgCl_2_, was washed out without noticeable effects on our preparations.

To test the possible role of different neurotransmitter candidates in cilia control, we applied them to the recording dish using a graduated 1 mL pipette attached to a long, small-diameter tube. The final concentrations were calculated from the known volume of injected solution and the known volume of the recording dish.

The following candidates for signal molecules were used in these experiments: GABA, acetylcholine, serotonin, glutamate, dopamine, histamine, glycine, aspartate, octopamine, FMRFamide, and two donors of nitric oxide (NOC-9 [6-(2-Hydroxy-1-methyl-2-nitrosohydrazino)-N-methyl-1-hexanamine], diethylamine NONOate [DEA NO or Diethylammonium (Z)-1-(N,N-diethylamino)diazen-1-ium-1,2-diolate], see details in (Maragos et al., 1991;Keefer et al., 1996;Braga et al., 2009;Li et al., 2020)). All chemicals were obtained from Sigma. Specifically, we used the following concentrations on both *Pleurobrachia bachei* and *Bolinopsis infundibulum*. Gamma-aminobutyric acid (GABA), total semi-intact preparations n=6, at concentrations: 0.1 mM, 0.2 mM and 0.5 mM (4 independent tests for each concentration– no effect); Acetylcholine (ACh), n=5, at concentrations: 0.1 mM, 0.2 mM, 0.5 mM and (2 independent tests for each concentration – no effect); Serotonin (5-HT), n=3, at concentrations: 0.1 mM and 0.5 mM (2 independent tests for each concentration – no effect); L-Glutamate, n=3, at concentrations: 2 times 0.5 mM and 0.2 mM (2 independent tests for each concentration – no effect); Dopamine (DA), n=3, at concentrations: 0.1 mM, 0.2 mM and 0.4 mM (3 independent tests for each concentration – no effect); L-DOPA (DA precursor); once at 0.5 mM – no effect; Histamine, n=3; at concentrations: 0.5 mM, 0.4 mM and 0.1 mM (3 independent tests for each concentration – no effect); Glycine, n=2; at concentrations: 0.4 mM, and 0.2 mM (2 independent tests for each concentration – no effect); L-Aspartate, n=2, 0.5 mM and 0.2 mM (2 independent tests for each concentration – no effect); Octopamine, n=2, 0.4 mM and 0.2 mM (2 independent tests for each concentration – no effect); FMRFamide, n=8 at concentrations: 0.2 mM and 0.1 mM (3-5 independent tests for each concentration – suppression of complex patterns of cilia activity in combs). Effects of NO donors: NOC-9, n=5, at 0.1 mM and 0.2 mM (3 independent tests for each concentration –inhibition of comb ‘s cilia beating; Diethylamine NONOate, n=13 at subsequent 0.02 mM, 0.06 mM, 0.1 mM, and 0.2 mM in seawater, and n=3 in high MgCl_2_ (3 independent tests for each condition – inhibition of comb ‘s cilia beating). Details about NO donors and FMRFamide are described in the result section.

To understand whether the possible effect was direct on the cilia cells or indirect via potential interneurons and due to chemical transmission, ‘chemical isolation ‘ was used by bathing the preparation in high Mg^2+^ saline for 5-15 minutes (333 mM MgCl_2_ was added to filtered seawater at a 1:1 ratio). Elevated magnesium chloride solution suppresses synaptic chemical transmission and is widely used in comparative neurobiology (Del Castillo and Engbaek, 1954;Hutter and Kostial, 1954). All solutions were prepared immediately before use. In all experiments, we checked the effect of a candidate neurotransmitter on the frequency of cilia beating and the occurrence and intensity of bursts. The cilia beating was compared before transmitter application, after application for about 5-30 minutes, and then after washing in seawater for about 5-15 minutes. The solution in the experimental chamber was changed within 5 mins by a five-fold volume of the fresh filtered (<0.2 µm) seawater.

Immunohistochemical labeling was performed as described elsewhere using anti-FMRFamide antibody (Cat # AB15348, Sigma-Aldrich). See details about the protocol and *Pleurobrachia* neuroanatomy (Norekian and Moroz, 2019;2020).

## RESULTS AND DISCUSSION

In semi-intact preparations, patterns of cilia beating in *Pleurobrachia* were variable, with periods of bursts and inhibitory episodes (**Figure 2**). Such activity might represent intact behaviors in free-moving *Pleurobrachia* as an ambush predator. In contrast, *Bolinopsis* had more regular, almost constant cilia beating with few activity patterns (**Figure 1C-E**), also reminiscent of its free-moving behavior.

**Figure 2.**
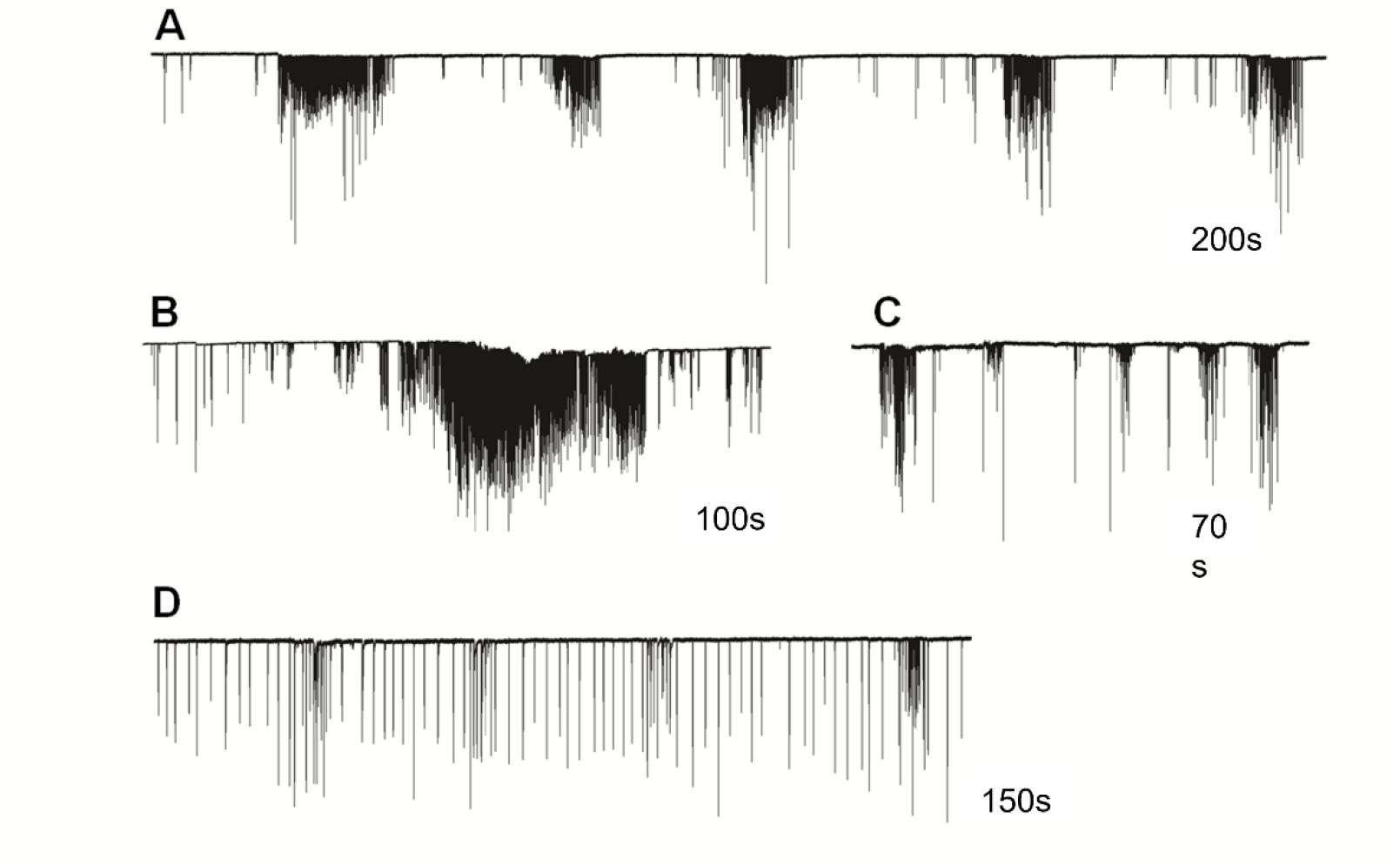
Cilia beating in *Pleurobrachia bachei* was very variable and complex, similar to intact behaviors in free-moving animals. For example, (A) regular episodes of high-frequency bursting with periods of inhibition between them, (B) long-duration powerful bursts of comb cilia strikes, (C) irregular unstructured bursting of cilia movements, (D) regular cilia beating with possible brief episodes of acceleration.

The irregular patterns of cilia activity were eliminated in the presence of a high concentration of Mg^2+^ (**Figure 3**), known to suppress synaptic inputs (Del Castillo and Engbaek, 1954;Hutter and Kostial, 1954). These findings indicate the presence of multifaceted regulatory chemical inputs and likely neuronal/secretory control of cilia, which was anticipated from ultrastructural data and neuro-ciliary synapses (Hernandez-Nicaise, 1991).

**Figure 3.**
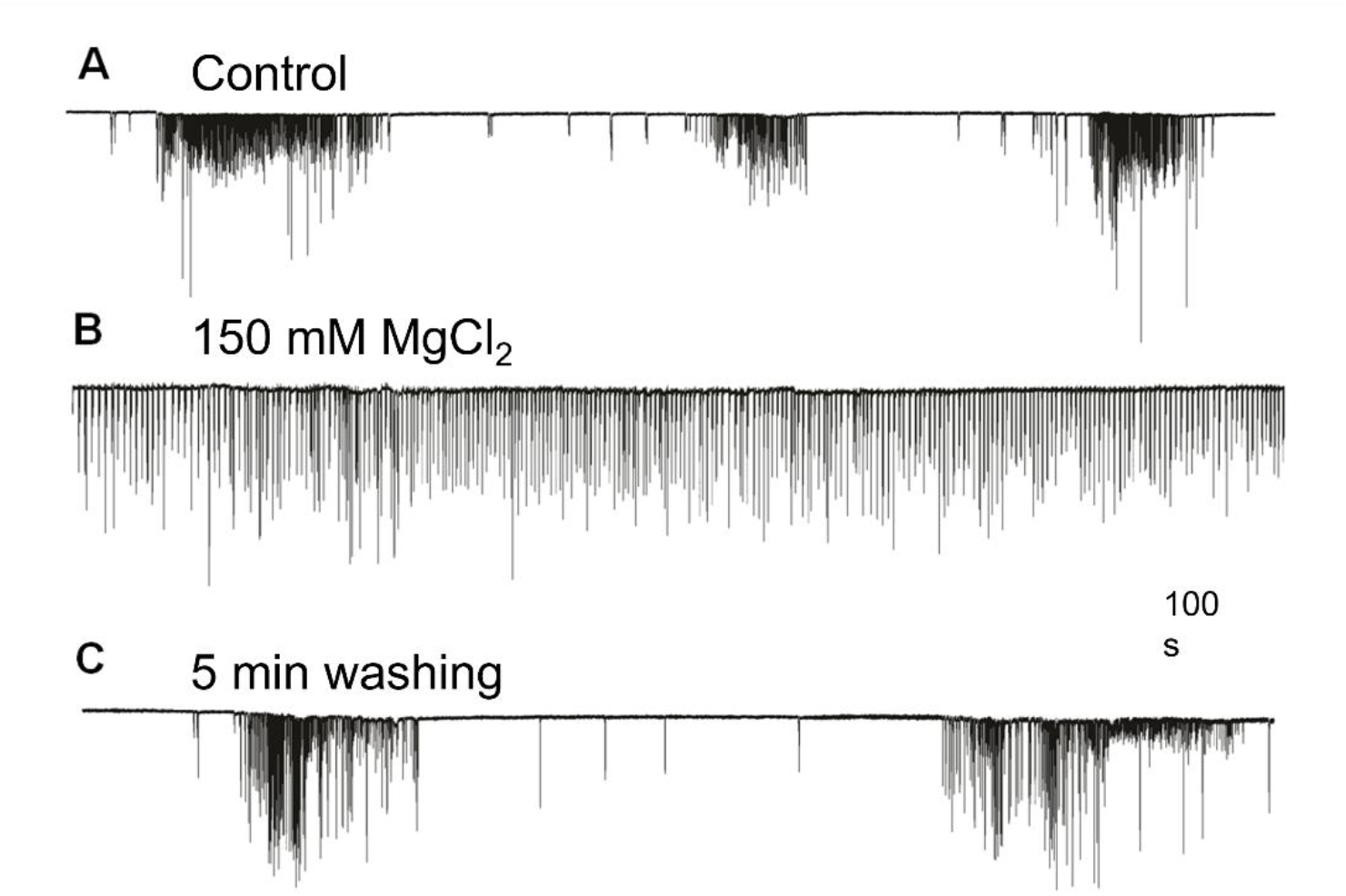
High Mg^2+^ seawater (B) blocked complex patterns of cilia ‘s beating (A), suggesting synaptic inputs initiated high-frequency bursting and inhibition episodes. The regular unvarying cilia beating in high Mg^2+^ solution (B) was removed by washing in normal seawater, fully restoring (C) previously observed episodes of bursting and inhibition. Numbers under all traces show the time of recordings.

Cilia excitatory and cilia inhibitory transmitters are unknown for most ctenophore species. A few neuropeptide candidates have been recently identified in *Mnemiopsis leidyi* (Sachkova et al., 2021) and *Bolinopsis* (Hayakawa et al., 2022) as candidate signal molecules controlling ciliated locomotion in these species. We performed pharmacological screening of low molecular weight transmitter candidates in semi-intact preparations. The effects of different signal molecules on *Pleurobrachia* cilia beating were similar to their actions produced in *Bolinopsis*: no observable effect on the application of selected low molecular weight transmitters and inhibitory action of nitric oxide donors (see below). Our screening showed an apparent lack of involvement of bilaterian neurotransmitters in the ctenophore cilia activity; Previous pharmacological and electrophysiological tests were consistent with the hypothesis that L-glutamate could be a neuromuscular transmitter in ctenophores because of its higher efficiency in inducing muscle contractions than D-glutamate and L-aspartate (Moroz et al., 2014). However, neither L-glutamate, L-aspartate, nor any other bilaterian amino acid-derived neurotransmitters tested here (glycine, GABA, acetylcholine, serotonin, dopamine, octopamine, histamine) could change the frequency of cilia beating in *Pleurobrachia* and *Bolinopsis* in concentrations up to 0.5 mM (see methods). These observations also support the hypothesis that acetylcholine and monoamines are bilaterian innovations (Moroz and Kohn, 2015;Moroz et al., 2021).

### Modeling Peptidergic Signaling

The first neural systems might have mainly been peptidergic (Moroz, 2009;2021). Peptidergic signaling can significantly affect interneuronal communication in ctenophores (Moroz et al., 2014;Sachkova et al., 2021;Hayakawa et al., 2022). *Pleurobrachia* and *Mnemiopsis* genomes do not encode FMRFamide (Moroz and Kohn, 2016). However, this versatile tetrapeptide might be used as a tool to mimic the action of some other endogenous short neuropeptides. Specifically, these peptides have multiple conformational states (Edison et al., 1999;Espinoza et al., 2000;Dossey et al., 2006) with affinity to various receptors because they are short. When the complete list of endogenous peptides is not determined precisely (as in ctenophores), RFamide related peptides can be efficiently used as a model for initial screening for the presence of peptidergic neurons and their actions. This approach was applied here as a part of screening for modulatory action on cilia activity in *Pleurobrachia*.

FMRFamide had an apparent inhibitory effect on high-frequency bursts of activity in cilia (**Figure 4**). In 10-20 seconds after application, the frequency of cilia beating in bursts was reduced, and the appearance of bursts was also decreased. The whole effect could be observed within 1-2 minutes. Of note, there was no effect of FMRFamide in high Mg^2+^ seawater (**Figure 5**, n=2). It suggests that the observed action of FMRFamide was indirect and polysynaptic.

**Figure 4.**
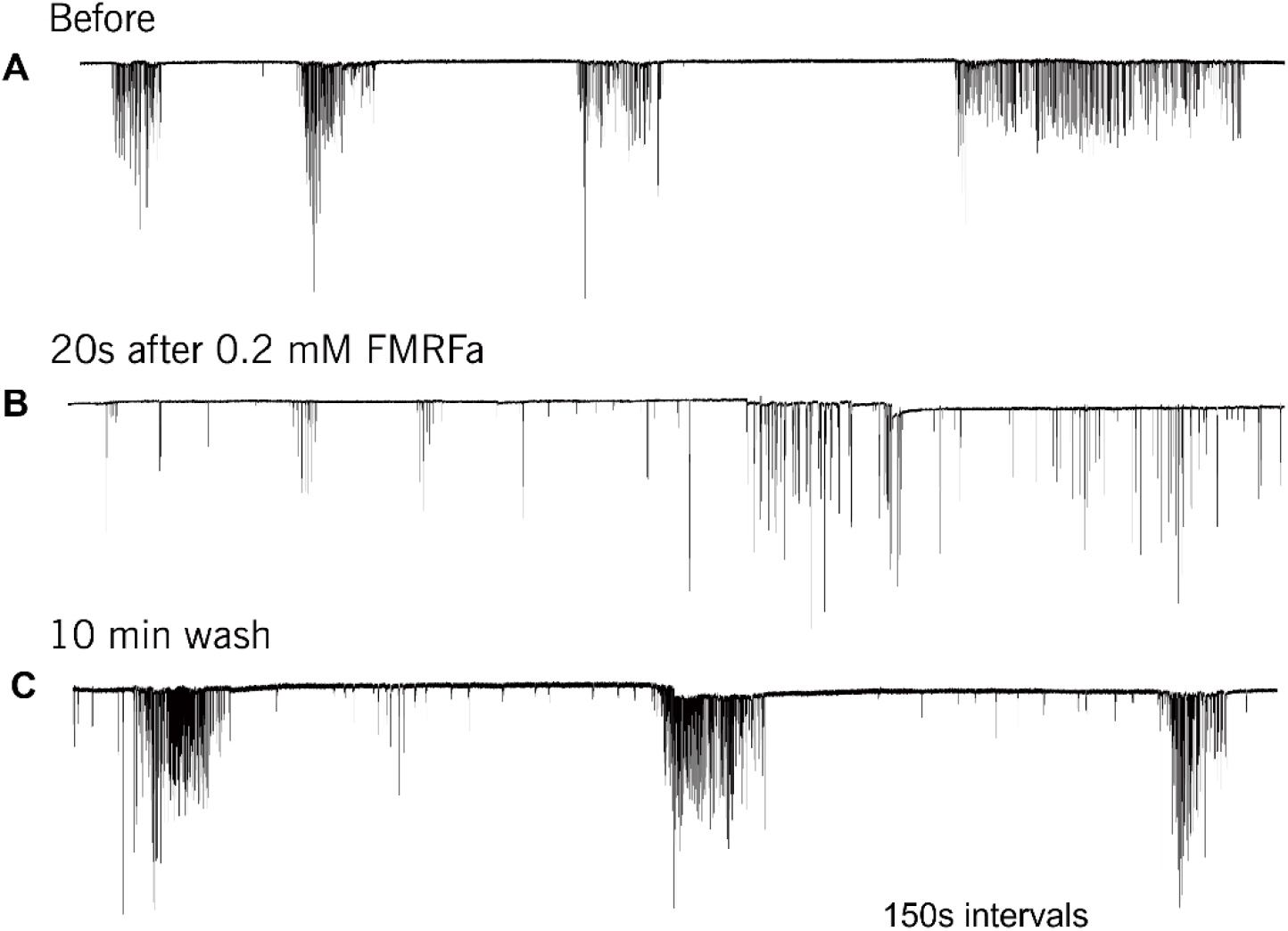
In *Pleurobrachia* FMRFamide reversibly inhibited the intensity of cilia bursting activity, repressing, or even eliminating the occurrence of bursts, significantly weakening the degree of cilia acceleration in the remaining bursts (A-C). Numbers under all traces show the time of recordings.

**Figure 5.**
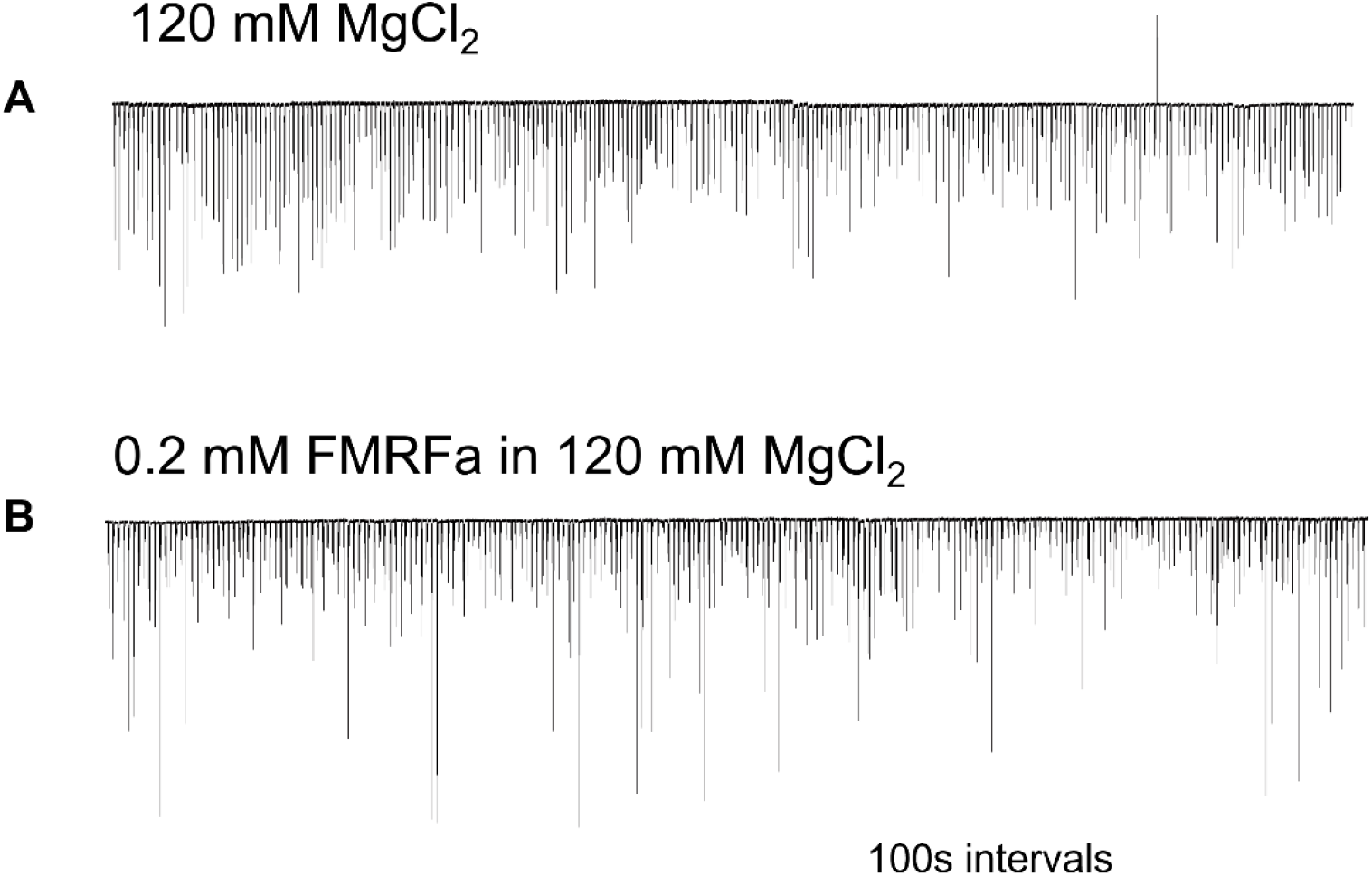
In high Mg^2+^ seawater (B), which blocked the variable bursting activity of *Pleurobrachia* cilia, FMRFamide did not demonstrate any noticeable effect on locomotory cilia, suggesting that its target was not comb ‘s cilia themselves but an external source controlling inputs to ciliated cells. Numbers under all traces show the time of recordings.

Because FMRFamide and other short peptides have rapid 3D confirmation dynamic in solutions (Edison et al., 1999;Espinoza et al., 2000;Dossey et al., 2006), we also assumed they might be cross-reactive with many endogenous peptides. We tested this situation using immunohistochemistry and revealed a distinct subset of peptidergic neurons, not reported previously. This is consistent with an observation that RFamide immunoreactivity was also detected in specific cells of the polar field in *Pleurobrachia* (**Figure 6**). These potentially chemoreceptive cells might use short neuropeptides as afferent components of neural circuits controlling locomotion via still known interneurons and motoneurons.

**Figure 6.**
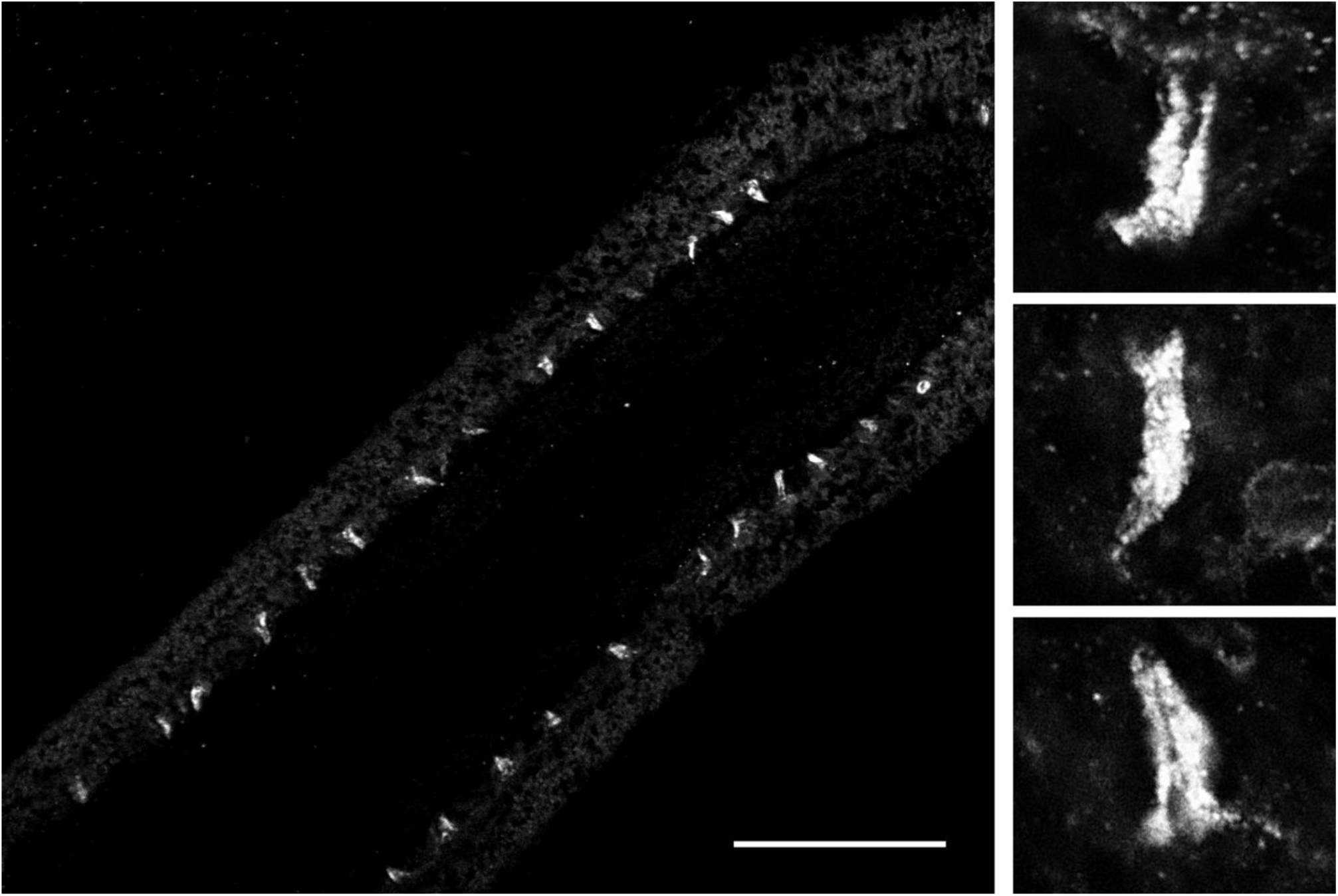
FMRF-like immunoreactivity in the polar field, a putative chemosensory organ of *Pleurobrachia bachei*. The left image shows regularly positioned stained putative chemoreceptive cells at lower magnification. Scale bar – 100 µm. Images on the right show the same cells at higher magnification. See details about the protocol and *Pleurobrachia* neuroanatomy (Norekian and Moroz, 2019;2020).

### Modeling Nitrergic Signaling

Nitric oxide (NO) is an ancient and versatile signal molecule (Moroz and Kohn, 2011a), recently proposed as a transmitter candidate in ctenophores (Moroz and Kohn, 2016;Moroz et al., 2023). In contrast to classical transmitters, the application NO donors (NOC-9 and Diethylamine NONOate, 0.02-0.2 mM) caused inhibition of comb cilia beating both in *Pleurobrachia* and *Bolinopsis* with a complete arrest of cilia activity in most cases at higher concentrations, >70-100 µm (**Figure 7**). The effect developed slowly over 2-5 minutes after NO-donor applications. This inhibitory effect was always reversible and was quickly washed out in the seawater with a complete restoration of pre-application activity in about 5 minutes. Of note, this inhibitory action of NO donors persisted in high Mg^2+^ seawater, suggesting the direct action of NO on the cilia in combs (**Figure 8**).

**Figure 7.**
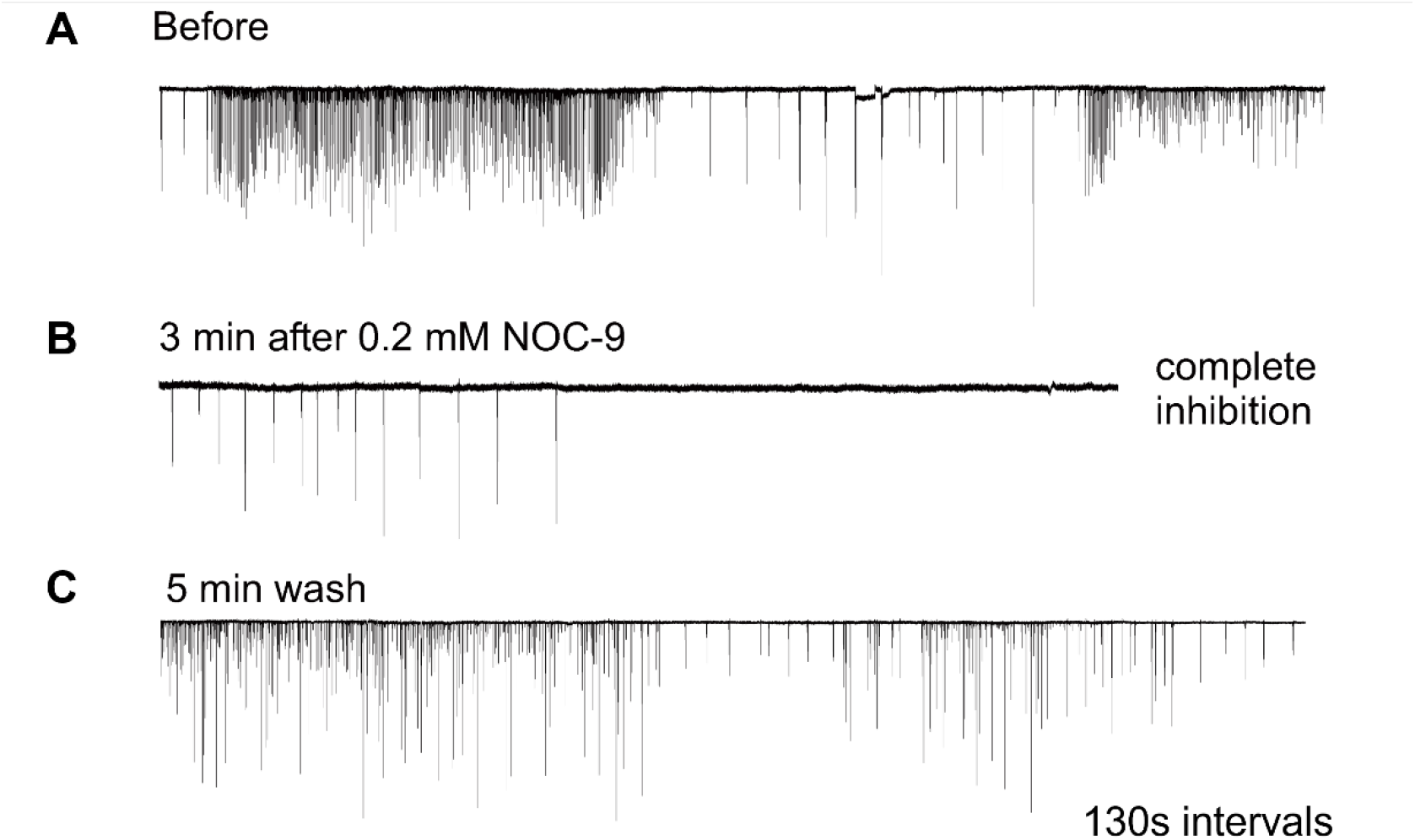
Nitric Oxide (NO) donors, such as NOC-9, reversibly suppressed cilia beating, completely inhibiting cilia movements in *Pleurobrachia* (A-B). This effect was reversible, and cilia beating was restored within 5 min of washing (C). Numbers under all traces show the time of recordings.

**Figure 8.**
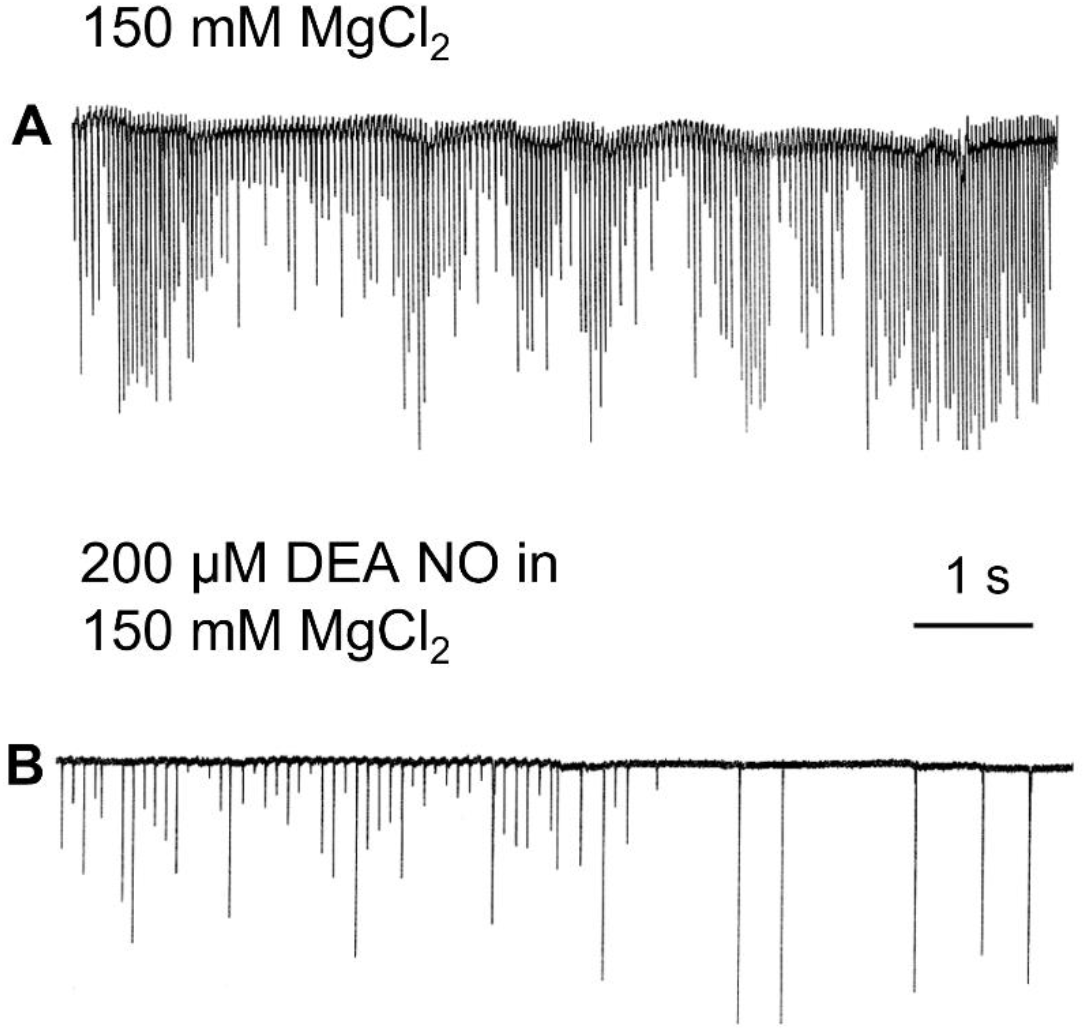
In high Mg^2+^ seawater, which blocked variable synaptic inputs and chemical transmission, Nitric Oxide (NO) donors suppressed cilia beating and even completely inhibited cilia movements in *Pleurobrachia* (A-B), suggesting that NO targets can be comb ‘s cilia themselves.

Our results imply that endogenous or environmental NO suppresses the cilia beating in ctenophores. Gaseous NO is one of the smallest and most diffusible signal molecules, with multiple non-enzymatic and enzymatic synthetic pathways, including nitric oxide synthase (NOS) in both host cells and microbiome (Moroz and Kohn, 2011b). Interestingly, the screening of the sequenced genome in *Pleurobrachia* and several transcriptomes from this species did not recover any NOS-like enzymes (Moroz et al., 2014). However, NOS was detected in basal and more derived species of ctenophores, such as *Mnemiopsis leidyi* (Moroz and Kohn, 2016;Moroz et al., 2020a) and *Bolinopsis* (Moroz et al., 2023). These comparative analyses illustrate the mosaic nature of NOS distribution within the phylum Ctenophora and provide evidence for the secondary loss of NOS in *Pleurobrachia* from the common ancestor of ctenophores (Moroz et al., 2023). However, *Pleurobrachia* has soluble guanylyl cyclases and possibly other receptosrs for NO, which might sense this molecule from alternative endogenous and exogenous sources (e.g., microbiomes and/or food).

## CONCLUSION & FUTURE DIRECTIONS

Ctenophores are one of the earliest, if not the earliest, lineage of metazoans (Whelan et al., 2015;Whelan et al., 2017;Li et al., 2021), central to understanding the origins and fundamental principles of animal organization. The life of ctenophores is entirely based on cilia, with dozens of populations of ciliated cells (Tamm, 1982;Hernandez-Nicaise, 1991;Tamm, 2014). As a result, multi-transmitter control of cilia activity is paramount to ctenophore organization and behaviors.

NO-cilia interactions can be one of the ancient signaling pathways in the evolution of animals, but this is in little investigated direction, with no comparative data (Saternos and AbouAlaiwi, 2018). Thus, it would be essential to identify both sources and mechanisms of the action of NO on cilia in different ecological groups of ctenophores. Experiments on other ctenophore species are imperative because of the mosaic distribution of NOS across species, with examples of secondary loss of this enzyme in many lineages (Moroz et al., 2023).

Second, the observed suppression of complex ciliary patterns by MgCl_2_ indicates the significance of steady state chemical transmission in generating ctenophore behaviors. This experiment is important because of the recently discovered syncytial organization of five ctenophore neurons in the subepithelial nerve net of early developmental stages of *Mnemiopsis* (Burkhardt et al., 2023). The finding might be interpreted as support for the widespread role of non-synaptic and non-chemical transmission in ctenophores (Dunn, 2023). However, the majority of neurons in ctenophores and external control of cilia activities are likely mediated by chemical transmission. Specifically, distinct ctenophore neural systems can employ well-recognized synapses already detected by electron microscopy (Hernandez-Nicaise, 1991;Burkhardt et al., 2023) and volume-type intercellular transmission (Moroz et al., 2021) mediated by small peptides, nitric oxide and, perhaps, additional low molecular weight messengers to be determined in future studies. The precise balance and complementary contributions of different transmitter mechanisms in ctenophores are the areas of exciting discoveries essential for fundamental neuroscience and deciphering the evolution of alternative integrative systems across basal metazoan lineages (Jekely, 2021;Moroz et al., 2021;Moroz and Romanova, 2022;Nikitin et al., 2023).

## AUTHORS CONTRIBUTIONS

TN and LLM designed the study and jointly performed experiments; TN and LLM wrote the manuscript; both authors reviewed and edited the manuscript.

## ACKNOWLEDGEMENTS

This work was supported in part by the Human Frontiers Science Program (RGP0060/2017) and National Science Foundation (IOS-1557923) grants to L.L.M. Research reported in this publication was also supported in part by the National Institute of Neurological Disorders and Stroke of the National Institutes of Health under Award Number R01NS114491 (to L.L.M). The content is solely the authors ‘ responsibility and does not necessarily represent the official views of the National Institutes of Health.

